# Hypogonadal (Gnrh1^hpg^) mice reveal niche-specific influence of reproductive axis and sex on intestinal microbial communities

**DOI:** 10.1101/2023.06.20.545808

**Authors:** Laura Sisk-Hackworth, Jada Brown, Lillian Sau, Andrew A. Levine, Lai Ying Ivy Tam, Aishwarya Ramesh, Reeya S. Shah, Evelyn T. Kelley-Thackray, Sophia Wang, Anita Nguyen, Scott T. Kelley, Varykina G. Thackray

## Abstract

The gut microbiome has been linked to many diseases with sex bias including autoimmune, metabolic, neurological, and reproductive disorders. Numerous studies report sex differences in fecal microbial communities, but how this differentiation occurs remains unclear. Using a genetic hypogonadal mouse model that does not produce sex steroids or go through puberty, we investigated how sex and the reproductive axis impact bacterial diversity within the small and large intestine. Both sex and reproductive axis inactivation altered bacterial composition in an intestinal section and niche-specific manner. Our results also implicated factors independent of the reproductive axis (i.e., sex chromosomes) in shaping intestinal communities. Additionally, our detailed profile of intestinal communities showed that fecal samples do not reflect bacterial diversity in the small intestine. Our results have ramifications for studying the impact of sex differences on the gut microbiome, particularly in sex-biased diseases and factoring in sex and steroid levels in microbial-based therapies.

## INTRODUCTION

Biological sex and the reproductive axis are fundamental to mammalian development and physiology. In addition to controlling sexual differentiation, fertility, and sex steroid production, the reproductive axis influences the nervous, musculoskeletal, cardiovascular, immune, hepatic, and gastrointestinal systems ^1-7^. Additionally, many diseases have a sex bias, including autoimmune disorders such as lupus, Crohn’s disease, and ulcerative colitis ^8,9^, or are influenced by sex steroids, such as polycystic ovary syndrome and cardiovascular disease ^5,10^.

Recently, sex and the reproductive axis have been implicated in regulating another critical aspect of mammalian biology, namely the gut microbiome. Numerous studies indicate that the human gut microbiome differentiates during puberty in a sex-dependent manner, coinciding with activation of the reproductive axis. Sex differences in the taxonomic composition of the gut microbiome (i.e., beta diversity) emerge during puberty, become more distinct in adulthood, and then diminish in older adults ^11-26^. A similar pattern occurs with alpha diversity, i.e., within-sample biodiversity, where sex differences are most pronounced in young and middle-aged adults but disappear in later adulthood ^19,20,22,23,25,27,28^. Sex differences in alpha and beta diversity have similarly been observed in rodents, indicating that mice make a good model for studying the connection between gut microbiota and sex-specific host physiology and diseases ^29-32^.

While there is growing evidence of a relationship between sex and the composition and diversity of the gut microbiome, the precise nature of this interaction is unclear. Thus far, published studies detecting sex differences in the gut microbiome have used fecal samples as a proxy for intestinal microbial communities. However, the intestinal tract has distinct physiological sections with unique abotic and biotic factors that create highly selective microhabitats for microbial communities, the diversity of which may not be captured by fecal samples. For example, low pH and high levels of bile acids and oxygen in the duodenum reduce microbial biomass and select for acid-tolerant, bile-acid transforming and oxygen-tolerant species, while in the cecum higher pH and lower oxygen levels select for anaerobic bacteria that ferment carbohydrates ^33-36^. Furthermore, intestinal epithelial cells secrete mucins and antimicrobials, creating distinct mucosal niches for mucin-degrading and mucin-adherent bacteria along the intestinal tract ^37-42^; for example, paneth cells produce a gradient of antimicrobial peptides that decreases from the proximal to distal small intestine ^33,43^. The mucosal barrier structure also shifts from the small to large intestine, with a thin mucus layer in the small intestine and a thick, two-layer mucosa in the cecum ^33^. The small intestine also has lower bacterial biomass and faster transit times than the large intestine ^33^. As evidenced by studies in humans, mice and other animals, these physiological differences select for distinct microbiomes that differ in microbial composition and biomass both lengthwise and cross-sectionally within the intestinal tract ^39,44-54^.

There is also evidence that the physiology of the intestinal tract is influenced by sex. For instance, one study showed that testosterone levels were high in the intestinal lumen of male mice, while estrogen levels were high in females, and these levels differed by intestinal site ^55^. Total bile acid levels within the intestine also been shown to vary by sex in humans and mice ^56-60^. Moreover, intestinal immune surveilance differs by sex. Schwerdtfeger and Tobet showed that mast cells were larger and more prevelent in females and paneth cells were more reactive in males when exposed to bacterial lipopolysaccharide ^61^. Collectively, these studies suggest that sex and the reproductive axis may play an important role in shaping the intestinal microbiome in a site-specific manner.

However, most studies that have focused on site-specific intestinal microbiota have not reported sex or have only assayed one sex ^44,47,53,54,62-67^, while studies that included both sexes did not test for sex differences ^34,39,42,50,68,69^. Recent studies have reported sex differences in the intestinal metabolome, but did not study the microbiota ^70^ or only tested for sex differences in the lumen of a specific intestinal section, but not the mucosa ^71,72^. Given the intense interest in the development of personalized microbial therapies for various human diseases ^73-75^, it is important to understand the influence of the reproductive axis and sex on microbial communities in both the small and large intestine.

In this study, we used the hypogonadal Gnrh1^hpg^ mouse model to test the hypothesis that sex and the reproductive axis shape niche-specific bacterial diversity of the gut microbiome. Hypogonadism in the *hpg* mouse stems from a large truncation of the *Gnrh1* gene, leading to a lack of gonadal development ^76,77^. In homozygous mutants, the reproductive axis is inactive, meaning that male and female *hpg*^*+/+*^ mice do not produce sex steroid hormones or go through puberty (Fig. 1A). In contrast to interventional methods such as gonadectomy, this model facilitates study of the effect of hypogonadism *starting in embryonic development* on maturation of the gut microbiome and isolates differentiation due to the hypothalamic-pituitary-gonadal (HPG) axis from other chromosomal-based sex differences. To investigate how ablation of the HPG axis alters sexual development of the intestinal microbiome, we assayed intestinal microbial communities from adult male and female wild-type and male and female *hpg* mutant mice (Fig. 1B). Since sex effects likely vary both radially and lengthwise along the intestinal tract, we performed 16S rRNA amplicon sequencing of luminal and mucosal samples from the duodenum, ileum, and cecum, as well as the feces of each mouse (Fig. 1C). Our results provide an unprecedented view into the impact of the reproductive axis on the mammalian gut microbiome, showing that the HPG axis drives development of intestinal sex differences and that the effect of sex and the HPG axis is specific to each intestinal niche.

**Fig. 1.**
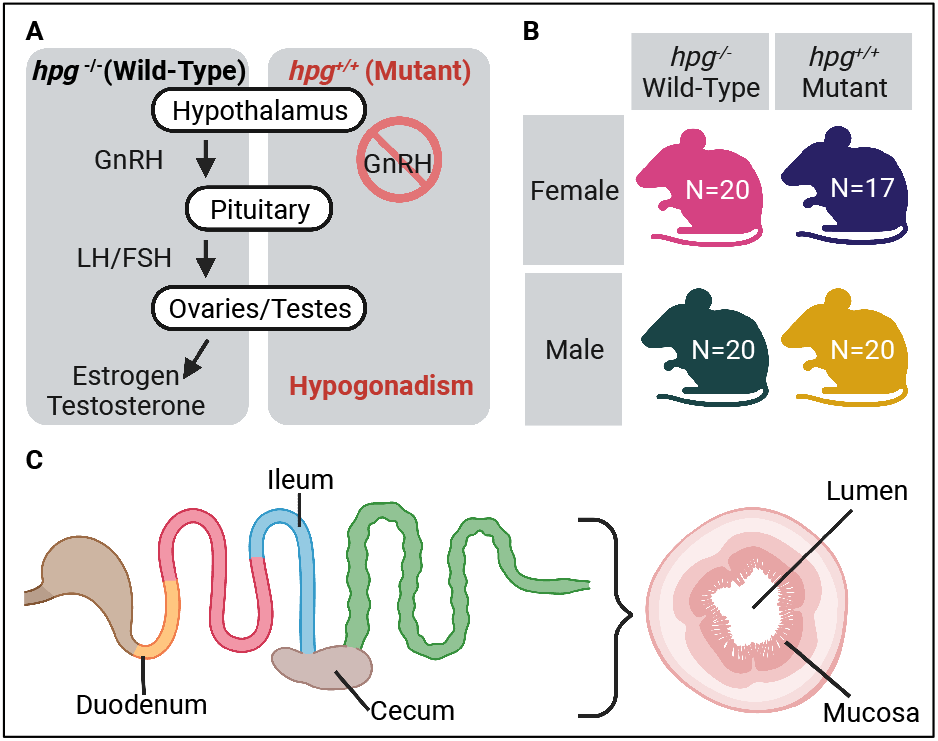
|Experimental Design. **A** In *hpg*^*-/-*^(wild-type) mice, the hypothalamic-pituitary-gonadal axis is functional. Gonadotropin-releasing hormone (GnRH) is secreted from the hypothalamus and stimulates production of luteinizing hormone (LH) and follicle-stimulating hormone (FSH) from gonadotrope cells in the anterior pituitary. LH and FSH then regulate steroidogenesis and gametogenesis in the ovaries and testes which results in sexual development and reproductive competence. In *hpg*^*+/+*^ mutant mice, GnRH is not produced due to a truncation in the *Gnrh1* gene, resulting in hypogonadism. **B** The study included four experimental groups: female and male wild-type and mutant mice. N = number of mice per group. **C** Sampling of intestinal microbial communities was done in the duodenum, ileum, and cecum, both in the lumen and mucosa, resulting in 539 samples including fecal samples.

## RESULTS

### Bacterial composition is driven by intestinal microhabitat, *hpg* genotype and sex

Beta diversity analysis of all samples combined found the strongest correlation of bacterial community composition with intestinal section (duodenum, ileum, cecum) or feces, followed by sample type (lumen vs. mucosa). NMDS ordination showed a clear separation between small intestine samples (duodenum, ileum) versus large intestine (cecum) and feces (Fig. 2A), a pattern confirmed by mixed-effects PERMANOVA tests (Section: p = 0.0001, R^2^ = 0.195; Supplemental Fig. 1). These results support the notion that each section harbors a unique microbial composition resulting from differential abiotic and biotic factors ^33,78^. Our data also detected significant differentiation between lumen and mucosa communities, supporting a role of the intestinal epithelium in shaping intestinal gut communities (Sample Type: p = 0.0001, R^2^ = 0.046; Supplemental Fig. 1). In addition, PERMANOVA identified an effect of both genotype and sex on microbial community composition, with significant interactions between sex and genotype, and genotype and section (Supplemental Table 1). Many of these distinctions were readily observed in the NMDS plots, such as the differentiation of ileum lumen and mucosa communities in female mutants that was not observed in female wild-type samples (Fig. 2A).

**Table 1:** R2 and p values from mixed-effect PERMANOVA for Euclidean distances. P values < 0.05 shown in bold.

| Fixed Effect | Duodenum |  | Ileum |  | Cecum |  |
| --- | --- | --- | --- | --- | --- | --- |
|  | P value | R2 | P value | R2 | P value | R2 |
| Sample Type | **0.0001** | 0.0571 | **0.0001** | 0.0260 | **0.0001** | 0.0147 |
| Sex | 0.1953 | 0.0070 | **0.0274** | 0.0098 | **0.0015** | 0.0103 |
| Genotype | **0.0047** | 0.0122 | **0.0007** | 0.0151 | **0.0001** | 0.0152 |
| Sex:Genotype | 0.1238 | 0.0076 | **0.0168** | 0.0102 | **0.0001** | 0.0119 |
| Sex:Sample Type | 0.0746 | 0.0083 | 0.9195 | 0.0047 | 1 | 0.0032 |
| Genotype:Sample Type | 0.8671 | 0.0048 | 0.3920 | 0.0063 | 1 | 0.0031 |
| Sex:Genotype:Sample Type | **0.0392** | 0.0091 | 0.2402 | 0.0070 | 0.9999 | 0.0032 |

**Fig. 2.**
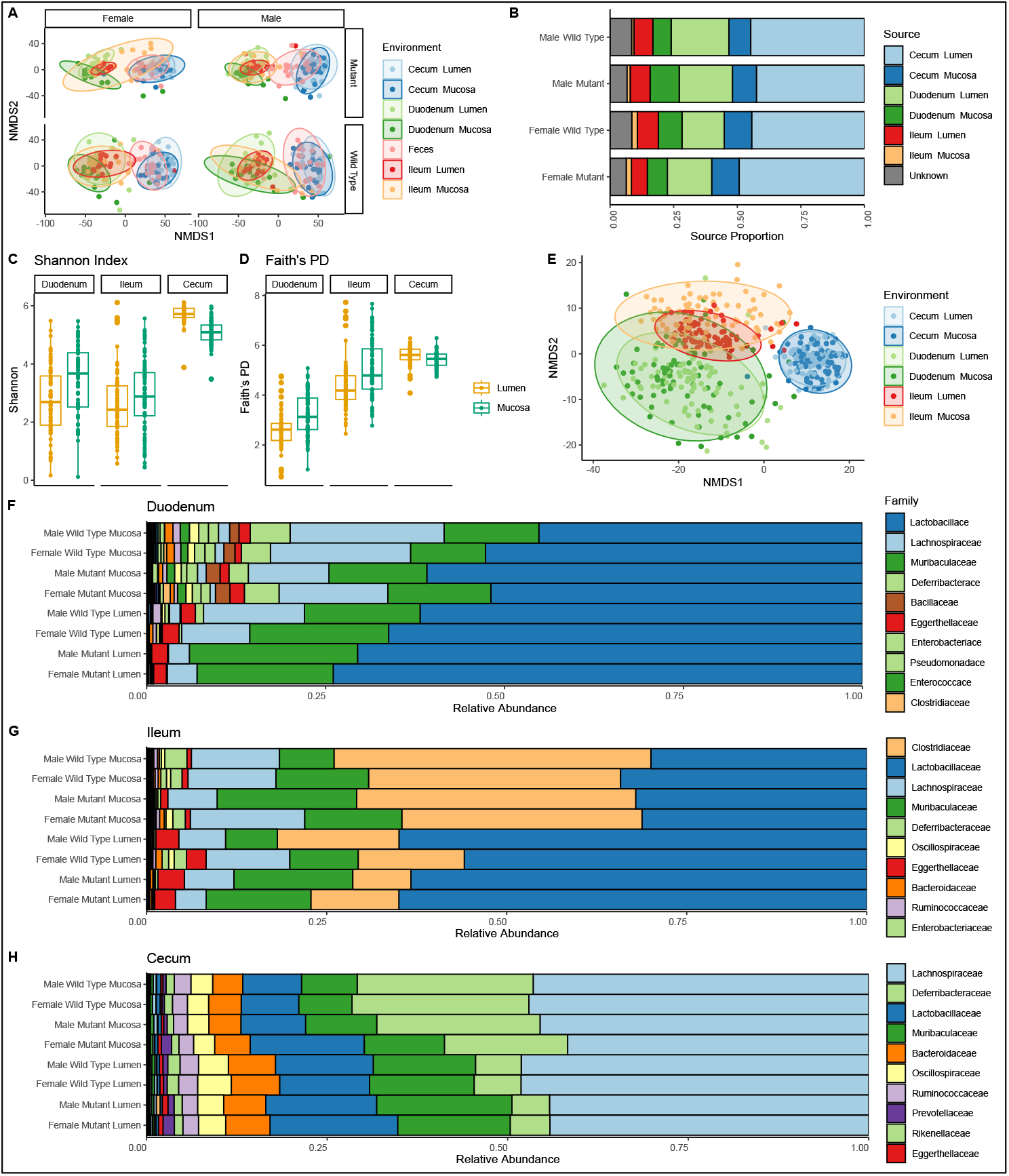
:Intestinal environment, sex, and *hpg* status affect microbial composition. **A** NMDS ordination of Euclidean distances determined from 16S sequence variant (SV) counts of samples collected from mouse intestinal environments (lumen and mucosa of duodenum, ileum, and cecum) and feces. Samples are colored by sample type and NMDS ordinations were split by sex and *hpg* genotype (Wild Type and Mutant). **B** Results of SourceTracker analysis using feces as sink and intestinal environments as sources. **C** Shannon Index and **D** Faith’s PD for the lumen and mucosa of duodenum, ileum, and cecum samples for all groups combined. **E** NMDS ordination of Euclidean distances based on bacterial family abundances for each intestinal environment. Bacterial family-level taxonomic bar plots for **F** Duodenum, **G** Ileum, and **H** Cecum samples.

### Fecal samples are not a good proxy for small intestinal communities

In all four experimental groups of mice, fecal samples consistently clustered with cecum samples and separately from small intestine (duodenum and ileum) samples (Fig. 2A). SourceTracker analysis also showed that the cecum lumen was the dominant source for the fecal communities (sink) and contributed 45% while cecum mucosa contributed 9.9% (Fig. 2B; Supplemental Fig. 1). Surprisingly, this was followed by the duodenum lumen, which contributed 28.5% while the duodenum mucosa contributed 9%. The ileum lumen contributed less than 10% and the ileum mucosa contributed negligibly to the fecal microbiome (Fig. 2B; Supplemental Fig. 1). Given that microbial composition in the intestine was not represented well in feces, we analyzed the effects of location, *hpg* genotype and sex on intestinal samples for the remainder of this study.

### Luminal and mucosal habitats influence alpha and beta diversity in a section-specific manner

Both intestinal section (duodenum, ileum, or cecum) and sample type (lumen or mucosa) had a strong effect on alpha diversity (Fig. 2C-D, Supplemental Table 2). The duodenum and ileum both had lower biodiversity at the SV (sequence variant) level (Shannon index; Fig. 2C, Supplemental Table 2), while overall phylogenetic diversity was lowest in the duodenum and highest in the cecum (Faith’s PD; Fig. 2D, Supplemental Table 2). Interestingly, in the duodenum and ileum, both Shannon and phylogenetic diversity were higher in mucosa than lumen, while the opposite was true in the cecum (Fig. 2C-D). There was no detectable correlation between alpha diversity and sex or *hpg* genotype in the duodenum or ileum (Supplemental Table 2). However, there was an interaction between alpha diversity, sex, and *hpg* genotype in the cecum (Faith’s PD; p = 0.017; Supplemental Table 2, Supplemental Fig. 2).

Differences in SV beta diversity between the lumen and mucosa were more apparent in the duodenum and ileum than in the cecum (Table 1, Supplemental Fig. 3) although PERMANOVA analysis showed significant differences between lumen and mucosa samples in all three sections for both Euclidean and UniFrac distances. In the duodenum, NMDS ordination of both abundance-based Euclidean and phylogenetic-based unweighted UniFrac distances showed a clear separation of lumen and mucosa samples (Supplemental Fig. 3). There was a greater difference in dispersion between ileum lumen and mucosa samples when accounting for abundances (Supplemental Fig. 3B), while the clustering of ileum lumen and mucosa samples was more distinct when accounting for phylogeny (Supplemental Fig. 3E). By contrast, in the cecum, NMDS ordination of Euclidean distances showed minimal differences between lumen and mucosa, and unweighted UniFrac distances showed mucosal samples were only slightly more dispersed than lumen samples (Supplemental Fig. 3C and F).

### Intestinal environment drives microbial composition at the bacterial family level

Beta diversity analysis at a higher taxonomic level showed clear differences in the community composition of the three intestinal sections at the family level, with the duodenum having the highest compositional variability while the cecum had the least (Fig. 2E). Furthermore, the duodenum and ileum communities were more alike than either were to cecum communities and, of the three sections, the ileum had the strongest differentiation between lumen and mucosa (Fig. 2E). Abundances of the top ten families in the intestine, *Lactobacillaceae, Lachnospiraceae, Muribaculaceae, Deferribacteraceae, Clostridiaceae, Bacteroidaceae, Oscillospiraceae, Eggerthellaceae, Ruminococcaceae*, and *Rikenellaceae* differed by section or sample type (lumen vs mucosa) or had a significant interaction between section and sample type (Supplemental Table 3).

Like family-level beta diversity, bacterial family relative abundance showed that duodenum and ileum communities were more similar to each other than to cecum communities. The lumen of both the duodenum and ileum had higher proportions of *Eggerthellaceae* and *Lactobacillaceae* compared to the cecum (Fig. 2F-H; Supplemental Fig. 4). However, the high proportion of *Clostridiaceae* in the ileum differentiated it from both the duodenum and cecum communities, and *Clostridiaceae* was higher in the ileum mucosa than the lumen (Fig. 2F-H). The duodenum was dominated by *Lactobacillaceae, Muribaculaceae, and Lachnospiraceae*, though *Lactobacillaceae* and *Muribaculaceae* were more abundant in the lumen, while *Lachnospiraceae* was more abundant in the mucosa (Fig. 2G). *Lachnospiraceae* dominated the cecum samples, which also had high relative abundances of several families that were in low abundance in the small intestine samples, including *Deferribacteraceae, Bacteroidaceae, Oscillospiraceae*, and *Ruminococcaceae* (Fig. 2H). Differences between lumen and mucosal communities in the cecum were also less distinct at the family level than in the small intestine sections (Fig. 2F-H). However, *Deferribacteraceae* was more relatively abundant in the cecum mucosa, while *Muribaculaceae, Lactobacillaceae, Bacteroidaceae, Oscillospiraceae, and Eggerthellaceae* were more abundant in the cecum lumen (Fig. 2H).

Differences in family-level diversity due to *hpg* genotype and sex were also apparent: *Bacteroidaceae, Eggerthellaceae, Muribaculaceae*, and *Rikenellaceae* were significantly different between wild-type and *hpg* mutant mice and *Muribaculaceae* was significantly different between sexes (Supplemental Table 3).

### Reproductive axis and sex niche-specifically impact intestinal community composition

Having detected relationships between microbial composition and sex or *hpg* genotype in the overall intestine (Supplemental Table 1), we performed a section-specific analysis of the impacts of these factors on the beta diversity of the duodenum, ileum, and cecum. Effects of *hpg* genotype on beta diversity were detectable in all sections with both Euclidean and unweighted UniFrac distances (Table 1; Supplemental Table 4). Given the significant effect of sample type, we performed NMDS ordination analysis separately for lumen and mucosa for each intestinal section. These analyses showed that *hpg* genotype altered community composition in both sexes within the lumen and mucosa of duodenum (Fig. 3A), ileum (Fig. 3C), and cecum (Fig. 3E), with *hpg* mutants consistently displaying less intra-sample variation than wild type in the lumen. Interestingly, samples from ileum mucosa showed the opposite pattern: NMDS ordination showed a higher degree of inter-sample variation in female *hpg* mutants than wild-type samples (Fig. 3C).

**Fig. 3.**
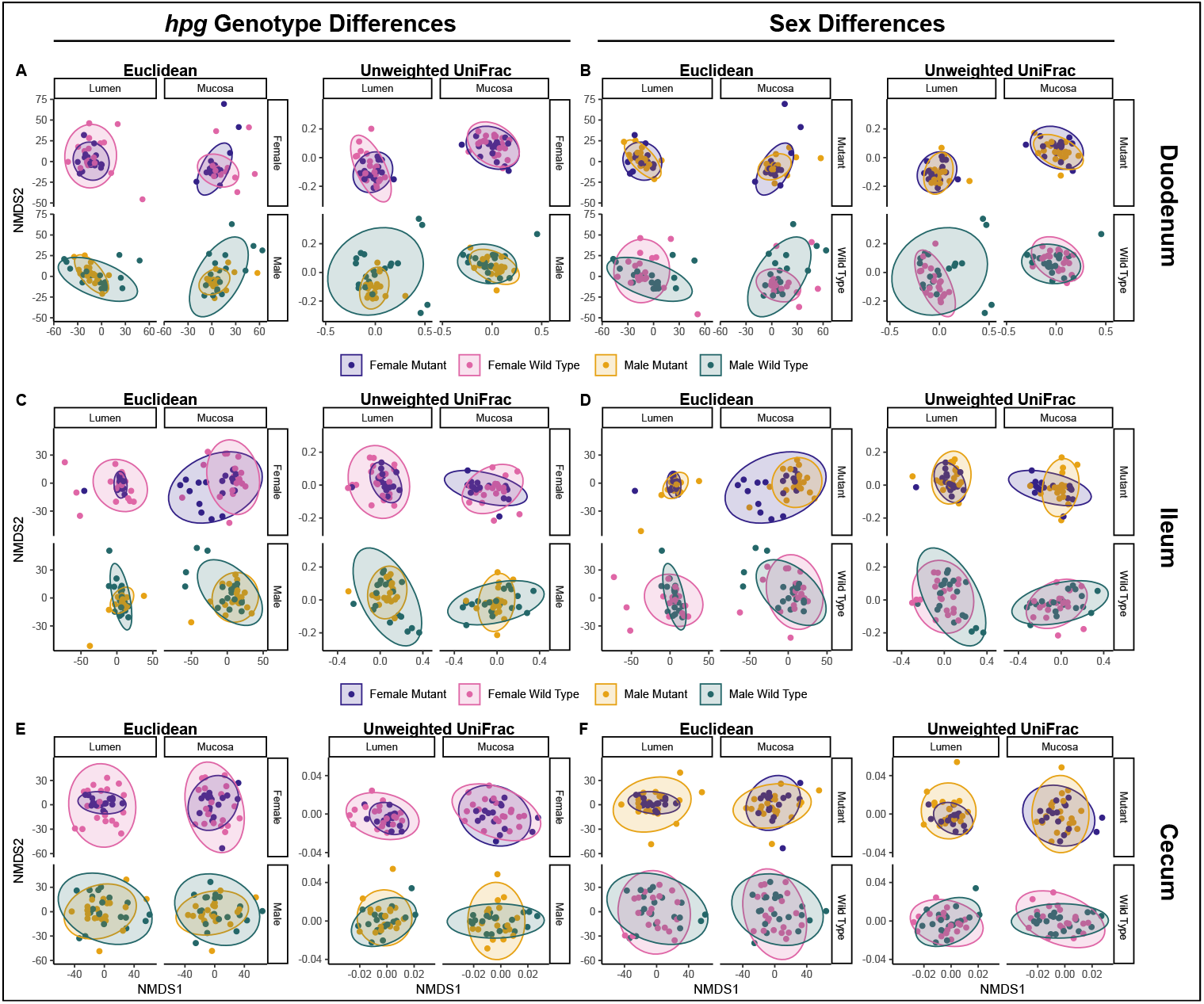
| Genotype and sex influence intestinal beta diversity in a niche-specific manner. Euclidean (clr-transformed SV abundance) and unweighted UniFrac (phylogenetic) distances revealed *hpg* genotype and sex differences in microbial communities of the small intestine and cecum. Genotype differences within wild-type and *hpg* mutant mice for **A** duodenal, **C** ileum and **E** cecal samples. Sex differences within male and female mice for **B** duodenal, **D** ileum and **F** cecum samples.

An effect of sex on beta diversity was also detectable in the ileum and cecum (Table 1; Supplemental Table 4). While sex did not affect beta diversity in the duodenum overall, there was a three-way interaction of sex, *hpg* genotype and sample type (Table 1). Differences between sexes in the duodenum were present in wild-type mice but not mutant mice and the dispersion differed by lumen and mucosa (Fig. 3B). In the ileum and cecum, sex correlated with microbial composition and there was an interaction between sex and genotype in the cecum (Table 1). In the ileum lumen, there were sex differences in wild-type but not *hpg* mutants, while in the mucosa, there were sex differences in both wild-type and mutant mice (Fig. 3D), indicating that chromosomal effects independent of the reproductive axis may also play a role in development of sex differences in the ileum. NMDS ordination also illustrated an increase in inter-sample variation of female mutants compared with male mutants in the ileum mucosa (Fig. 3D). In the cecum, sex differences in composition were less pronounced than in the ileum (Fig. 3F).

Additionally, the effect of the HPG axis in the cecum was similar for males and females, with NMDS ordination showing greater inter-sample variation among wild-type compared to mutant mice (Fig. 3F).

### Niche-specific differentiation of bacterial abundances by sex and *hpg* genotype

Given that the HPG axis and sex had distinct effects on overall beta diversity in the duodenum, ileum, and cecum, we then used the compositional algorithm, coda4microbiome to determine groups of genera whose abundances relative to each other collectively differentiate samples in each intestinal section microbiome by sex and genotype. Figure 4 shows the microbial balances that defined sex differences in both wild-type and mutant mice in addition to *hpg* genotype differences in female and male mice in the duodenum, ileum, and cecum for genera contributing most strongly to the balance (Fig. 4) or for all genera in the balance (Supplemental Fig. 5). We found that the wild-type sex difference balance model performed better (had a higher area-under-the-curve (AUC) score) compared to the mutant model in the ileum. In addition, the wild-type sex difference balance model was more parsimonious (indicating a stronger sex-specific signal) compared to mutant mice in the duodenum and cecum (Fig. 4; Supplemental Fig. 5). We also found that balance models comparing wild-type and mutant mice performed better (had a higher AUC value) for female mice than for male mice, indicating that the differences between wild-type and mutant females were more consistent (Supplemental Fig. 5). The models also showed that distinct bacterial genera contributed to the balances that defined sex differences in wild-type mice versus mutant mice and that some of these genera also contributed to genotype difference in female or males (Fig. 4D). For example, *Lachnoclostridum* in the duodenum contributed to the balance defining females versus males in wild-type mice and also contributed to the balance defining wild-type versus mutants in females (Fig. 4 A,D). Since the lumen and mucosa samples were combined for the balance analysis, heatmaps showed that the relative abundance of genera identified in the balance analysis differed in wild-type males and females between lumen and mucosa, particularly in the duodenum and ileum (Supplemental Fig. 6 A-C).

**Fig. 4.**
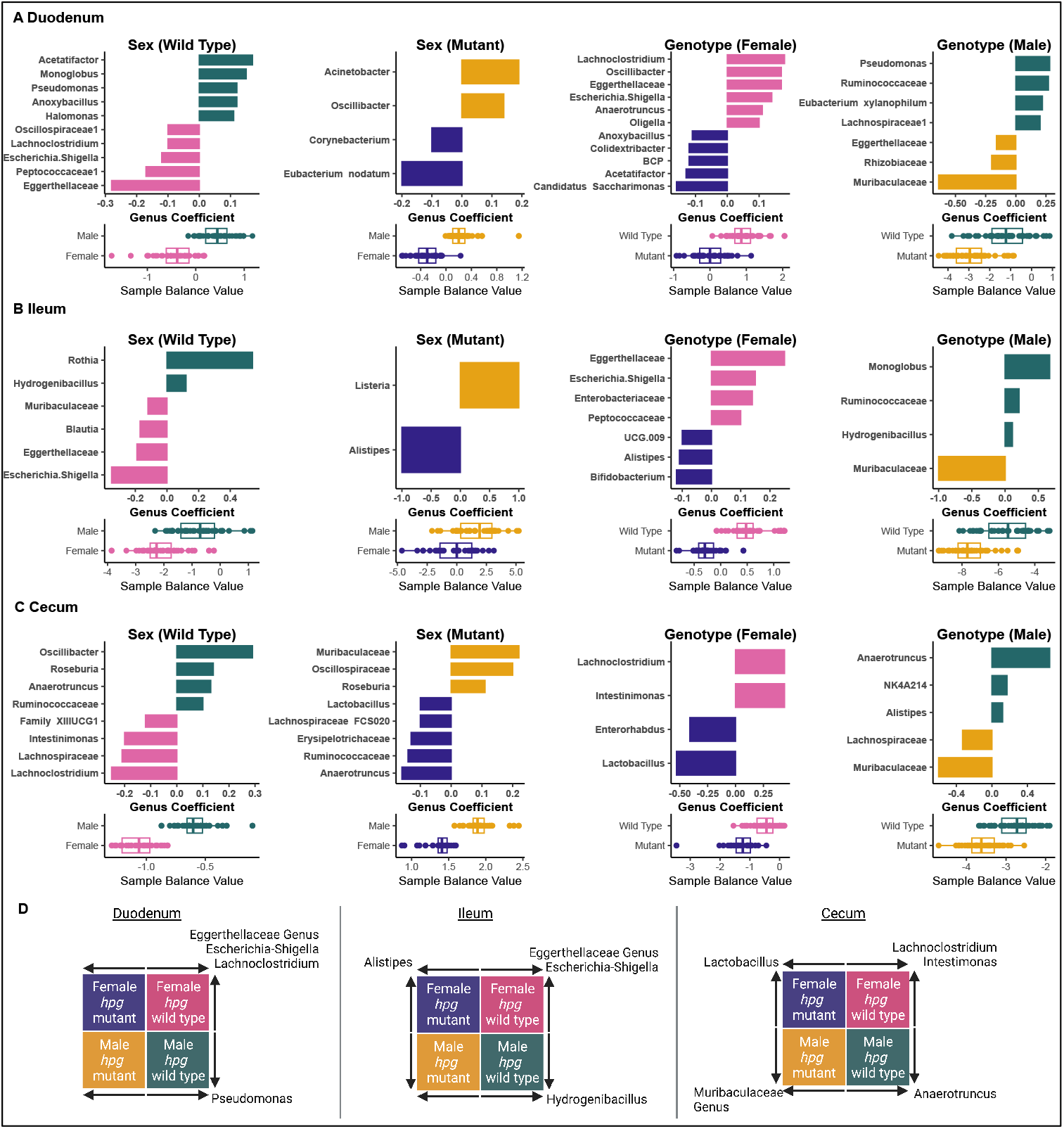
| Genus-level bacterial balances defined intestinal community differences by sex and genotype. Genus-level microbial balances defining wild-type sex differences, mutant sex differences, female *hpg* genotype differences, and male *hpg* genotype differences are shown for **A** duodenum, **B** ileum, and **C** cecum. For each balance, coefficients of genera contributing to the balance are shown with bar graphs. The length of the bar (Genus Coefficient) indicates the proportion that each genus contributed to the balance; genera contributing an absolute balance of at least 0.05 are shown. The direction of the bar indicates whether the genus contributed positively or negatively to the balance, and indicates in which group (male, female, mutant or wild-type) the genus was more abundant. Balance values for each sample in the specified groups are shown below each coefficient graph. For instance, in A, *Acetatifactor* contributed positively to the balance, indicating *Acetatifactor* had a higher relatively abundant in male than female wild-type samples. **D** Schematic showing genera with a coefficient of at least 0.05 that contributed to both sex and genotype differences, with the arrow indicating the group in which the genus was relatively more abundant within each balance. Abbreviation: BCP: *Burkholderia/Caballeronia/Paraburkholderi* genus group.

**Fig. 5.**
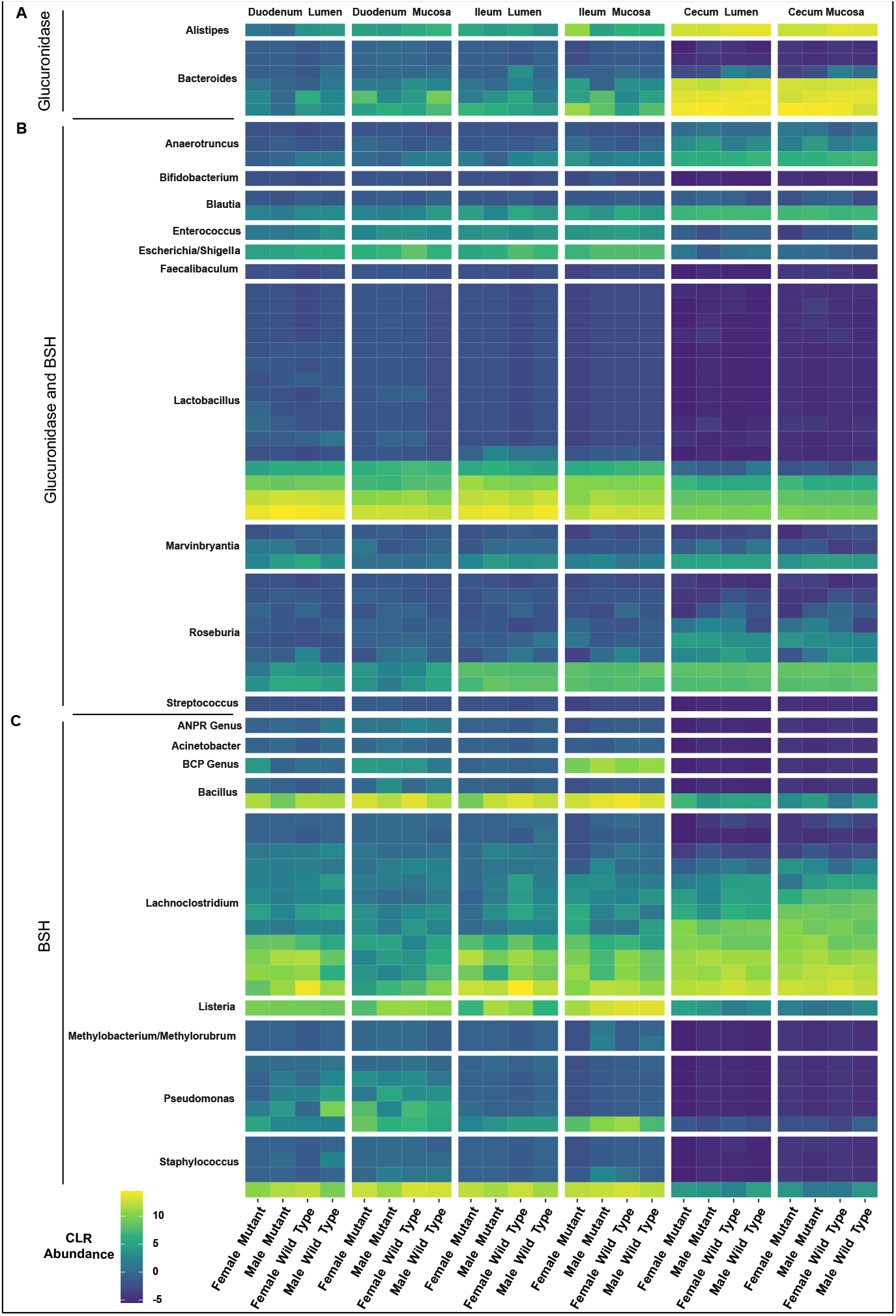
| Clr-transformed abundances of SVs belonging to genera with known glucuronidase and BSH activity. **A** Glucuronidase, **B** glucuronidase and bile-salt hydrolyase (BSH), and **C** BSH. Abbreviations: ANPR genus: *Allorhizobium/Neorhizobium /Pararhizobium/Rhizobium* genus group. BCP genus: *Burkholderia/Caballeronia/Paraburkholderi* genus group.

### Abundances of SVs with putative glucuronidases and bile-salt hydrolases are intestinal section-specific

Given that levels of sex steroids and bile acids differ between males and females and between intestinal environments, we next asked how the abundances of SVs belonging to bacterial genera with species known to have glucuronidases (an enzyme that can deconjugate steroid glucuronides) or bile salt hydrolases (BSHs) differ by intestinal environment, *hpg* genotype, and sex. For example, numerous species of *Lactobacillus* produce glucuronidases, BSHs or both ^79-81^. The heatmap in Figure 5 shows the mean clr-transformed counts for all SVs belonging to these genera separated by lumen and mucosa within intestinal section. *Bacteroides* and *Alistipes* SVs, which were known to have activity or the gene for glucuronidase but not BSH, were generally more prevalent in the cecum than the small intestine (Fig. 5A). Genera with both known glucuronidase and BSH genes or activity did not follow a clear pattern in intestinal abundance (Fig. 5B). SVs from genera with known BSH genes or activity were generally more abundant in the small intestine, besides *Lachnoclostridium*, which was more abundant in the cecum (Fig. 5C). In addition, the abundance of SVs with putative glucuronidases or BSHs differed by *hpg* genotype and sex. Notably, SVs within each genus did not follow the same abundance patterns between intestinal sections, showing how grouping SVs at the genus level can obscure abundance patterns at lower taxonomic levels (Fig. 5).

## DISCUSSION

Studies in humans and rodent models have shown that the fecal microbiomes of biological males and females are similar prior to puberty but then diverge in a sex-dependent manner during puberty. This pattern strongly suggests that activation of the HPG axis is responsible for sex differences in the gut microbiome but how this axis affects the microbial communities along the intestinal tract is unknown. In humans, it is very difficult to investigate the influence of the reproductive axis *per se* on the gut microbiome or to distinguish its effects from other sex chromosome-influenced factors. Furthermore, direct sampling of microflora along the human intestinal tract is a considerable challenge as is controlling for diet, which is a major confounding factor in gut microbiome studies.

Thus, our use of a genetic mouse model that knocks out *Gnrh1* gene expression in embryonic development and prevents activation of the HPG axis provides a powerful means to isolate the effects of the reproductive axis and readily examine its impacts on the microbiomes in different intestinal sections (both lumen and mucosa), while controlling for the effects of diet. In this study, we showed for the first time that the reproductive axis and sex exert niche-specific effects on intestinal microbial communities, with pronounced effects on overall bacterial community composition (beta diversity) and abundances of specific bacteria at the family, genus, and SV level.

In comparing mouse intestinal and fecal microbiomes, we found that fecal samples poorly represented the bacterial composition of the duodenum and ileum, though the cecum was relatively well represented. This mirrored previous studies, which found that fecal samples were more like samples from the large intestine than the small intestine ^36,46,47,54,82^. SourceTracker analysis also found that the strongest, most consistent source for fecal microbes was the cecum and, while there was a signal from the duodenum, the ileum was barely represented.

Thus, if the impacts of a condition or treatment on the microbiome are most pronounced in the small intestine, they may be difficult or impossible to detect in fecal samples. Differential taxonomic abundances in the duodenum could easily be masked or even contradicted by data from cecum communities which contain orders of magnitude more bacteria. This could explain why fecal studies have not identified sex differences in members of the *Eggerthellaceae* family while our study found sex differentiation in the relative abundances of *Eggerthellaceae* in the small intestine.

Our comprehensive taxonomic mapping of mucosal and luminal communities of the duodenum, ileum, and cecum found an extremely strong relationship between intestinal environment and microbial community diversity and composition. In agreement with prior studies ^46,47^, we found that the small intestine had lower alpha diversity than the cecum, with the lowest alpha diversity in the duodenum. This may be a consequence of the duodenum’s higher oxygenation, lower pH, and high transit times compared with other intestinal sections, resulting in greater environmental stochasticity. Greater environmental stochasticity tends to reduce community alpha diversity over time ^83^; however, stochastic events in an environment can create temporal niches that allow low-abundance taxa or otherwise poor competitors to survive ^83,84^. For example, the presence of *Halomonas*, which are aerobic or facultatively anaerobic, in the duodenum may be due to sudden increases in the oxygen level after eating due to the stomach depositing oxygenated contents into the duodenum or due to increased blood flow to the epithelial cells after food consumption ^85,86^. High environmental stochasticity may also explain why the duodenum had the greatest inter-individual community variability at the family level and the highest number of low abundance SVs. In the small intestine, mucosal communities had higher levels of alpha diversity than luminal communities, while in the cecum the differences in alpha diversity between mucosa and lumen were small or nonexistent. This suggests that the mucosa may protect against the low pH and bile acids in the duodenum and ileum that make survival challenging to many taxa ^87,88^.

Differences in intestinal environments were also reflected in bacterial family-level community composition. In agreement with prior studies, members of the acid-tolerant, bile acid-transforming *Lactobacillaceae* ^87,89^ were a much higher proportion of small intestine communities than the cecum, and were also a higher proportion of luminal than mucosal communities in the small intestine ^34,47,54,90^. In addition to *Lactobacillaceae*, other abundant families in the duodenum included *Muribaculaceae, Lachnospiraceae*, and *Deferribacteraceae. Muribaculaceae* (previously named S24-7) has been positively associated with intestinal short-chain fatty acid (SCFA) levels and possesses genes for bile acid deconjugation ^91-93^. *Lachnospiraceae* and *Deferribacteraceae* species have been identified to have host mucin-degrading activity and activate immune signaling ^40,94^. The duodenal mucosa also had a higher relative abundance of *Bacillaceae, Enterobacteriaceae, Pseudomonadaceae, Staphylococcaceae, Enterococcaceae*, and *Oscillospiraceae* compared to the lumen, all of which have previously been associated with intestinal mucosa communities or mucin-degrading activity ^95-97^. Only two studies have compared the duodenal lumen to the duodenal mucosa in mice ^44,98^, so these findings add to the limited prior knowledge about the mucosal community in the duodenum.

While *Muribaculaceae* and *Lachnospiraceae* were also relatively abundant in the ileum, the most striking finding in the ileum was the high proportion of *Clostridiaceae*. The most abundant member of *Clostridiaceae* in the ileum was *Candidatus Arthromitus*, an uncultured segmented filamentous bacteria known to attach primarily to ileal epithelial cells ^99,100^. Another interesting finding was the relatively high proportion of *Eggerthellaceae* in the small intestine.

Although commonly isolated from human and animal feces ^96,101,102^, little is known about the role of *Eggerthellaceae* species in the gut, though they have genes for host mucin degradation ^96^. Additionally, our findings agreed with another study^39^ that *Lachnospiraceae* dominate the cecum and that the cecum mucosa has a higher relative abundance of *Deferribacteraceae* compared to the lumen; however, we also present new evidence that the cecum lumen maintains higher relative abundances of *Muribaculaceae* and *Lachnospiraceae* compared to the mucosa.

Using the *hpg* mouse model, we also identified several bacterial families whose relative abundances were influenced by the reproductive axis. The relative abundances of *Bacteroidaceae, Eggerthellaceae, Muribaculaceae*, and *Rikenellaceae* were significantly different between *hpg* and wild-type mice, and this effect varied between luminal and mucosal communities and by section. Since members of these families are known mucin degraders ^96,103,104^, one possibility is that differences between *hpg* mutant and wild-type host mucin production resulted in altered abundances of these families. Additionally, members of *Eggerthellaceae* and *Bacteroidaceae* have genes for bile acid metabolism ^105^ and members of *Bacteroides* and *Rikenellaceae* have putative glucuronidase activity ^106^, so bile acid levels and sex steroid levels may influence the competitive advantages of these taxa within the intestinal tract.

The effects of the HPG axis and sex were even more noticeable at the SV level. We showed that the effect of the reproductive axis and sex was more pronounced on overall composition (beta diversity) of intestinal microbial communities than biodiversity (alpha diversity). Beta diversity was strongly influenced by *hpg* genotype in each of the three intestinal sections, though the degree of differentiation was section- and site-(lumen vs. mucosa) specific. For example, genotype differences between wild-type and *hpg* mutant beta diversity in the duodenum were more pronounced in the lumen than the mucosa. Sex differences were also detectable in each intestinal section, though again the effects were section- and site-specific. For example, only wild-type mice showed sex differences in the duodenum, while in the ileum both wild-type and mutant *hpg* mice showed sex differences, though the later was only apparent in ileum mucosa samples. Collectively, these results strongly support the hypothesis that activation of the reproductive axis in wild-type mice due to *Gnrh1* gene expression results in a differentiation of intestinal microbial communities in adults that does not occur in *hpg* mutant mice. These results also indicate that the effects of sex and the reproductive axis are highly differential within the intestinal tract.

One of the most consistent patterns across all intestinal sections was a difference in SV-level community composition between wild-type mice and *hpg* mice, with NMDS plots showing greater dispersion in wild-type mice of both sexes compared to *hpg* mice. This result indicates that a principal effect of the HPG axis on the microbiome is an increase in inter-individual bacterial diversity. This agrees with prior findings showing greater inter-individual diversity in fecal samples between adult humans than between pre-pubertal children ^107^. It is possible that this occurs because activation of the reproductive axis increases habitat complexity within the intestinal tract due to its effects on the host (e.g., sex steroids, bile acids, immune system ^44,108-112^). Habitat complexity tends to increase beta diversity and niche differentiation ^113-115^. This finding also suggests that intestinal microbial communities in *hpg* mutant mice may be constrained to pre-pubertal-like levels of habitat complexity, resulting in lower levels of inter-individual variability.

In addition to overall bacterial composition, we found that the microbial balances defining sex and *hpg* genotype differences were niche-specific. Interestingly, some genera characteristic of differences between wild-type male and females also differentiated wild-type mice from mutants of the same sex. For example, in the duodenum *Lachnoclostridium* and *Escherichia Shigella* were more abundant in the balances differentiating wild-type females from wild-type males *and* wild-type females from mutant females. Similar patterns were observed in the ileum and cecum although the bacteria did not completely overlap. This pattern is interesting, as a metabolic feature common to members of *Lachnoclostridium* and *Escherichia Shigella* is the ability to deconjugate bile acids, which are at higher levels in female mice ^56,116,117^. Some of the other genera that were part of microbial balances for wild-type sex differences also have putative glucuronidase or BSH activity. For example, *Anaerotrunctus* (glucuronidase and BSH activity) and *Pseudomonas* (BSH activity) drove part of the balance for wild-type males in the duodenum and cecum, respectively, and *Lachnoclostridium* (BSH activity), *Blautia* (glucuronidase and BSH activity), and *Escherishia Shigella* (glucuronidase and BSH activity) tipped the microbial balance towards wild-type females in different parts of intestine ^79-81,106^. In addition, our results showed that SVs with putative glucuronidase and/or BSH activity did not have the same abundance patterns within different sections of the intestinal tract, and abundances were influenced by *hpg* genotype and sex. These findings highlight the potential for bile acids and sex steroids to regulate intestinal microbial communities and the need for future mechanistic studies to understand how these biotic factors mediate sex differences in intestinal communities.

Since the reproductive axis was embryonically inactivated in *hpg* mice, we were also able to show that host sex differences not related to gonadal sex steroids also shaped the gut microbiome in a section-specific manner. For instance, the sex differences we observed in SV-level community composition between *hpg* mutant male and females in the ileum mucosa and in genus-level balances cannot be a result of sex steroids because *hpg* mutant mice do not produce gonadal steroid hormones. This indicates that another mechanism, like sex chromosomal effects, led to sex differences in these instances. This finding opens a fundamental new area of investigation focused on genetic effects of sex chromosomes not mediated by the HPG axis on the gut microbiome. These effects might include the *SRY* gene or copy number of X chromosomes, which have been shown to influence physiology independently of sex steroids^118^.

While studies have compared luminal and mucosal communities of the small intestine or cecum in mice and humans, few have compared the duodenum lumen and mucosa ^47,98^, none have explored sex differences in small intestine lumen and mucosa, and none have investigated the influence of the reproductive axis on differentiation of intestinal microbial communities. Our study indicates the importance of including both sexes in gut microbiome studies, since disease or treatment effects in males may not correspond to effects in females and vice versa. We also demonstrate the importance of employing intestinal samples when studying sex differences, and emphasize that simply because a sex difference is not observed in fecal samples it cannot be assumed that there are no sex differences in intestinal microbial communities. In addition, the effects of gonadal steroid insufficiency/excess and treatments such as hormone replacement therapy on gut health should be further investigated. Moreover, for diseases with a sex bias that have a gut microbiome connection, sex differences in the gut microbiome should be explored as a mechanism for the sexual dimorphism. Furthermore, sex and hormonal status may need to be considered for personalized microbial-based therapies and fecal microbiota transplants. Finally, future studies are needed to investigate the mechanisms through which the HPG axis regulates differentiation of intestinal microbiomes and how these mechanisms relate to human health and disease.

## METHODS

### *Hpg* Mouse Model

The hypogonadal (*hpg*) mice were purchased from Jackson Labs (RRID:IMSR_JAX:000804). The *Gnrh1*^*hpg*^ mutation is a ∼33.5 kb deletion that removes two exons that encode most of the protein. Since *hpg* homozygous mice are infertile, *hpg* heterozygous male and female mice were crossed with each other and the offspring were genotyped to identify *hpg*^+/+^ (mutant) or *hpg*^-/-^(wild-type) homozygous males and females. Two cohorts of breeding mice were generated with the aim of producing 20 mice per genotype and sex. In the final tally, cohort A consisted of 8 female mutants, 10 female wild types, 11 male mutants, and 11 male wild types, while cohort B consisted of 9 female mutants, 10 female wild types, 9 male mutants, and 9 male wild types. Mice were housed in a vivarium with a 12-hour light (6:00 AM to 6:00 PM) and 12-hour dark cycle and had access to water and food (Teklad Global 18% Protein Extruded Diet; Envigo, Indianapolis, IN) *ad libitum*. Mice were single housed to prevent coprophagy from influencing the gut microbiome. The University of California, San Diego Institutional Animal Care and Use Committee approved all animal procedures used in this study (Protocol Number S14011).

### Sample Collection and DNA Isolation

When mice were 10 weeks of age, mice were weighed and fecal samples were collected, then mice were sacrificed using 2.5% isoflurane and cervical dislocation. The duodenum (first 7 cm of the intestine), ileum (last 7 cm before the cecum), and cecum were excised from the intestine. Luminal contents were collected by squeezing the contents of each section into a tube. Once contents were collected, the intestinal mucosa were gently shaken in three washes of PBS to remove remaining lumen material. Samples were frozen immediately after collection (luminal contents) or washing (lining mucosa) and stored at −80°C. To minimize potential variability induced by the estrus cycle, female wild-type samples were collected during estrus which was determined by vaginal epithelial cell smears. Bacterial DNA was extracted from the fecal and intestinal samples and from two negative extraction controls and two ZymoBIOMICS Microbial Community Standard positive extraction controls (D6300 Zymo Research) with the DNeasy PowerSoil Pro Kit (Qiagen) according to the manufacturer’s instructions, and stored at −80°C.

### 16S rRNA Amplicon Sequencing

The V4 hypervariable region of the 16S rRNA gene was PCR amplified using “universal” bacterial primers 515F and 806R, with the 806R primers barcoded with unique 12-bp Golay barcodes ^119^. Amplifications were performed separately for each environment (feces, cecum, ileum, and duodenum), with each set including two negative extraction controls, two negative PCR controls, two DNA extraction positive controls (ZymoBIOMICS Microbial Community Standard), and two PCR positive controls (ZymoBIOMICS Microbial Community DNA Standard). The feces and cecum samples were amplified using the following steps: an initial denaturation temperature of 94°C for 3 minutes, then 25 cycles of 45 seconds denaturation, 60 seconds of 50°C annealing, 90 seconds of 72°C extension, then a 72°C final extension for ten minutes. For the ileum and duodenum samples, the number of numbers of cycles was increased to 35 due to the low bacterial biomass, the annealing temperature was increased to 59°C to decrease potential amplification of host mitochondrial ribosomal RNA, and the extension time was decreased to 30 seconds to reduce amplification of spurious bands greater than 300 base pairs in length (the size of the bacterial V4 region).

Amplicon sequencing libraries were prepared by The Scripps Research Institute Next Generation Sequencing Core. Four sets of amplicons were used in separate library construction and sequencing runs with: (1) 77 fecal samples, (2) 154 combined cecum lumen and mucosa samples, (3) 154 ileum combined lumen and mucosa samples, and (4) 154 duodenum combined lumen and mucosa samples. Each run also included two positive and two negative extraction controls, and two positive and two negative PCR controls. Amplicon products were cleaned with Zymo DNA Clean & Concentrator™-25 columns, quantified using a Qubit Flourometer (Life Technologies), and pooled.

Sequencing libraries were prepared with the recommended Illumina protocol involving end-repair, A-tailing and adapter ligation. For the Ileum and duodenum, the prepared DNA library was size-selected on a 2% agarose gel (410–470 bp) using the Agencourt SPRI system to reduce mitochondrial contamination (Beckman Coulter, Inc.). After library prep, libraries were PCR amplified with HiFi Polymerase (Kapa Biosystems) for 12 cycles. Quantitation, denaturation in 0.1 N NaOH, then dilution to 5pM of libraries preceded loading libraries onto an Illumina single read flow-cell for sequencing on the Illumina MiSeq.

### 16S rRNA gene sequence quality control and QIIME2

Raw sequences (forward reads only) were processed with QIIME 2 (version 2022.8) ^120^. Reads and metadata were imported with qiime tools import. Reads were demultiplexed using the QIIME 2 cutadapt plugin ^121^. This resulted in 8.58 million total reads with 34,616 average reads per sample for the fecal run, 8.64 million total reads with an average 51,439 reads per sample for the cecum run, 8.10 million total reads with an average 48,279 reads per sample for the ileum run, and 6.71 million total reads with an average 39,946 reads per sample for the duodenum run. Observed sequence variants (SVs) were produced with QIIME 2 plugin dada2, truncating at 240 base pairs based on quality scores ^122^.

SV taxonomy was assigned with QIIME 2’s feature-classifier plugin using a pretrained naïve Bayes classifier trained on reference database Silva 138 ^123-125^. Zymo positive sequencing controls were evaluated for expected and unexpected sequences. The percent genus abundance matched closely to expected genera abundance, while unexpected sequences (potential contaminants), comprised less than 1% of total SVs in the positive controls.

Negative controls had low sequence counts and had between 39 and 672 times fewer counts than positive controls, and the composition of negative controls differed greatly from experimental samples. SVs belonging to non-gut bacteria families that were likely environmental contaminants (e.g., Deinococcaceae, Thermaceae, Thermoanaerobacterales Family III, Geodermatophilaceae, and Chitinophagaceae) were removed using qiime taxa filter-table. These potential contaminant families were not present in the cecal or fecal experimental samples and were only present at 0.099% and 0.012% of total SVs in the duodenum and ileum experimental samples, respectively. Sequences classified as mitochondria were also removed with qiime taxa filter-table. Additional mitochondrial SVs were identified by blasting taxonomically “unassigned” SV sequences against the *Mus musculus* mitochondrial 12S sequence and removed from the SV table.

Removal of likely contaminants resulted in 741 SVs in the fecal samples, 606 SVs in the cecum samples, 832 SVs in ileal samples, and 1554 SVs in duodenum samples. These SVs were then zero-filtered to remove very low-abundance SVs using the CurvCut heuristic approach^126^, which suggested feature removal of SVs present in 3 or fewer samples, resulting in a final counts of 333 fecal SVs, 309 cecum SVs, 286 ileal SVs, and 281 duodenum SVs.

Filtered ASV taxonomy was collapsed to the species, genus, and family levels using the QIIME2 plugin qiime taxa collapse. A rooted tree was constructed from the SVs using QIIME 2 plugin phylogeny align-to-tree-mafft-fasttree ^127,128^. Shannon Index ^129^, Faith’s Phylogenetic Diversity (PD) ^130^, and UniFrac distances ^131,132^, were calculated using command qiime diversity core-metrics-phylogenetic. Rarefaction depth was based on where alpha diversity metrics leveled off, excluding 2 low-sequence count samples (mouse 461 duodenum mucosa and mouse 414 cecal mucosa).

### Statistical Analysis

Analysis of SVs and diversity metrics were performed in R version 4.2.1. For family relative abundance bar plots, SVs were summed by family and family counts were made relative to the total sample counts. All graphs were generated using ggplot2 3.4.0. ^133^. SV, genus, and family count tables were transformed with the centered-log ratio to account for the compositional nature of the data ^134,135^. Euclidean distances for SV and family data were computed using clr-transformed feature tables with R vegan package 2.6.4 ^136,137^. NMDS ordination and PERMANOVA of Euclidean distances and unweighted UniFrac distances were also performed using vegan. Linear mixed-effect models were performed with package nlme version 3.1.161 ^138^ and FDR corrected for multiple comparison. Fixed effects were multiplied and random effects were mouse nested within cohort.

Microbial balances, coefficients, and AUC values were calculated with coda4microbiome ^139^. The coda4microbiome approach, an update to the selbal program ^140^, identifies the most parsimonious microbial balances defining differences between microbial communities using penalized logistic regression ^139^. coda4microbiome outputs a log-contrast model for the balances between two different microbial communities; positive microbial coefficients indicate that the microbe contributes to the community with a higher balance, while negative microbial coefficients indicate that the microbe contributes to the community with a lower balance. The absolute value of the coefficient indicates the relative amount that the specific microbe contributes to the balance model. To generate heatmaps showing the abundances of glucuronidase and bile salt hydrolase (BSH) functions, we identified bacterial genera from the literature that included species with demonstrated glucuronidase activity and/or BSH activity, or whose genomes encoded glucuronidase or BSH genes ^79,80,106^. Microbial source tracking of fecal samples was done with SourceTracker 2 using the diagnostics function on clr-transformed SV count tables ^141,142^. Confidence intervals were calculated in Python version 3.10.

## DATA AVAILABILITY

Sequence reads from this study were deposited into SRA under BioProject PRJNA983444. The QIIME 2, SourceTracker, Python, and R code used to analyze the data, as well as the taxonomic features tables (SV-, family-, and genus-level) and metadata with barcodes are available at (https://github.com/laurasisk/hpg_axis_intestinal_microbiome).

## ACKNOWLEDGEMENTS

This work was supported by NIH R01 HD095412 to V.G.T and S.T.K. L.S-H. was funded by NIH F31 HD105403, the Rees Stealy Research Foundation and the San Diego Chapter of the ARCS Foundation.

