## Supplemental Data for "Hypogonadal (Gnrh1^hpg^) mice reveal niche-specific influence of reproductive axis and sex on intestinal microbial communities"

This supplemental data file contains:
Supplemental Tables 1-4
Supplemental Figures 1-6

| Fixed Effect | p value | Df | SumOfSqs | R2 | F |
|---|---|---|---|---|---|
| **Section** | **0.0001** | 2 | 464289 | 0.1953 | 70.5253 |
| **Sample Type** | **0.0001** | 2 | 110964 | 0.0467 | 16.8554 |
| **Sex** | **0.0134** | 1 | 8017 | 0.0034 | 2.4355 |
| **Genotype** | **0.0004** | 1 | 14102 | 0.0060 | 4.2840 |
| **Sex:Genotype** | **0.0065** | 1 | 9119 | 0.0038 | 2.7702 |
| **Sex:Sample Type** | 0.5466 | 2 | 5900 | 0.0025 | 0.8962 |
| **Genotype:Sample Type** | 0.8430 | 2 | 4946 | 0.0021 | 0.7514 |
| **Sex:Section** | 0.2070 | 2 | 7614 | 0.0032 | 1.1566 |
| **Genotype:Section** | **0.0473** | 2 | 10197 | 0.0043 | 1.5489 |
| **Sample Type:Section** | **0.0002** | 2 | 27763 | 0.0117 | 4.2172 |
| **Sex:Genotype:Sample Type** | 0.2931 | 2 | 7019 | 0.0030 | 1.0662 |
| **Sex:Genotype:Section** | 0.2066 | 2 | 7542 | 0.0032 | 1.1457 |
| **Sex:Sample Type:Section** | 0.6726 | 2 | 5424 | 0.0023 | 0.8240 |
| **Genotype:Sample Type:Section** | 0.8259 | 2 | 4993 | 0.0021 | 0.7585 |
| **Sex:Genotype:Sample Type:Section** | 0.2861 | 2 | 7064 | 0.0030 | 1.0730 |
| **Residual** | | 511 | 1682031 | 0.7076 | |
| **Total** | | 538 | 2376984 | 1 | |

**Supplemental Table 1 | Location, sex and *hpg* genotype affect compositional differences between samples.** P values, degrees of freedom (DF), sum of squares (SumOFSqs), R2, and F statistics from mixed-effect model PERMANOVA for effects of section, sample type (lumen vs. mucosa), sex, and *hpg* genotype. P values < 0.05 shown in bold.

| Fixed Effect | Shannon Index | | | | | |
|---|---|---|---|---|---|---|
| | **Duodenum p value** | **Duodenum Chisq** | **Ileum p value** | **Ileum Chisq** | **Cecum p value** | **Cecum Chisq** |
| **Sample Type** | **6.63E-08** | 29.1692 | **0.0173** | 5.6614 | **2.20E-39** | 172.4077 |
| **Sex** | 0.9971 | 1.25E-05 | 0.1686 | 1.8948 | 0.1890 | 1.7247 |
| **Genotype** | 0.1288 | 2.3062 | 0.4723 | 0.5164 | 0.5405 | 0.3744 |
| **Sex:Genotype** | 0.6619 | 0.1911 | 0.5185 | 0.4167 | 0.7815 | 0.0769 |
| **Sex:Sample Type** | 0.5894 | 0.2911 | 0.5485 | 0.3599 | 0.8356 | 0.0430 |
| **Genotype:Sample Type** | 0.8476 | 0.0368 | 0.3929 | 0.7298 | 0.0537 | 3.7214 |
| **Sex:Genotype:Sample Type** | 0.2929 | 1.1060 | 0.3646 | 0.8217 | 0.5487 | 0.3596 |
| **Fixed Effect** | **Faith's PD** | | | | | |
| | **Duodenum p value** | **Duodenum Chisq** | **Ileum p value** | **Ileum Chisq** | **Cecum p value** | **Cecum Chisq** |
| **Sample Type** | **1.43E-09** | 36.6251 | **7.90E-08** | 28.8294 | **6.71E-07** | 24.6965 |
| **Sex** | 0.8486 | 0.0364 | 0.0506 | 3.8190 | 0.5933 | 0.2851 |
| **Genotype** | 0.1600 | 1.9737 | 0.8487 | 0.0363 | 0.9787 | 0.0007 |
| **Sex:Genotype** | 0.6575 | 0.1964 | 0.6192 | 0.2469 | **0.0487** | 3.8835 |
| **Sex:Sample Type** | 0.3540 | 0.8589 | 0.9291 | 0.0079 | **0.0387** | 4.2739 |
| **Genotype:Sample Type** | 0.4950 | 0.4655 | 0.1119 | 2.5258 | 0.3934 | 0.7284 |
| **Sex:Genotype:Sample Type** | 0.4467 | 0.5789 | 0.1322 | 2.2659 | **0.0169** | 5.6968 |

**Supplemental Table 2 | Sex and hpg genotype affect alpha diversity in the cecum only**. P value and Chi-square (Chisq) shown for linear mixed-effect model to determine effects of sex, *hpg* genotype, and sample type (lumen vs. mucosa). P values < 0.05 shown in bold.

| Fixed Effect | Bacteroidaceae | Clostridiaceae | Deferribacteraceae | Eggerthellaceae | Lachnospiraceae |
|---|---|---|---|---|---|
| **Sample Type** | **1.158E-04** | 1.053E-01 | **1.067E-06** | **1.447E-25** | **2.709E-03** |
| **Section** | **1.068E-79** | **0.000E+00** | **3.456E-50** | **1.279E-22** | **6.520E-36** |
| **Genotype** | 0.7974 | 0.5403 | 0.3247 | **0.0033** | 0.7974 |
| **Sex*Genotype** | 0.7119 | 0.7475 | 0.9501 | 0.7119 | 0.7119 |
| **Genotype*Section** | **0.0056** | 0.5062 | 0.5062 | 0.3226 | 0.6400 |
| **Sample Type*Section** | **1.267E-03** | **1.267E-03** | 1.271E-01 | **2.005E-08** | **7.914E-04** |
| **Sex*Genotype*Sample Type*Section** | 0.4384 | 0.2403 | 0.2403 | 0.0712 | 0.0715 |

| Fixed Effect | Lactobacillaceae | Muribaculaceae | Oscillospiraceae | Rikenellaceae | Ruminococcaceae |
|---|---|---|---|---|---|
| **Sample Type** | **2.504E-53** | **8.330E-30** | 1.870E-01 | **1.695E-03** | 0.1382 |
| **Section** | **2.472E-173** | **5.422E-22** | **6.587E-39** | **9.833E-114** | **4.912E-27** |
| **Genotype** | 0.1031 | **1.627E-05** | 0.9377 | 0.5142 | 0.5142 |
| **Sex*Genotype** | 0.8381 | **0.0401** | 0.7119 | 0.7119 | 0.7119 |
| **Genotype*Section** | 0.6400 | **0.0056** | 0.9392 | **0.0056** | 0.6400 |
| **Sample Type* Section** | **2.602E-19** | **8.596E-15** | **4.180E-02** | **1.480E-02** | **1.942E-03** |
| **Sex*Genotype*Sample Type*Section** | 0.0712 | **0.0124** | 0.1690 | 0.2403 | 0.2403 |

**Supplemental Table 3 | Effect of section, sample type, sex, or *hpg* genotype on top 10 most abundant bacterial families clr-transformed abundances.** FDR-corrected p values shown for linear mixed-effect model. Fixed effects were section*sample type*sex**hpg*. Non-significant interactions were removed. P values < 0.05 shown in bold.

| Fixed Effect | Duodenum | | Ileum | | Cecum | |
|---|---|---|---|---|---|---|
| | P value | R2 | P value | R2 | P value | R2 |
| **Sample Type** | **0.0001** | 0.0811 | **0.0001** | 0.0512 | **0.0014** | 0.0173 |
| **Sex** | 0.1687 | 0.0079 | **0.0198** | 0.0150 | 0.0745 | 0.0103 |
| **Genotype** | **0.0016** | 0.0175 | **0.0257** | 0.0139 | **0.0182** | 0.0122 |
| **Sex:Genotype** | 0.1587 | 0.0080 | 0.1146 | 0.0093 | **0.0105** | 0.0134 |
| **Sex:Sample Type** | 0.0723 | 0.0095 | 0.9800 | 0.0023 | 0.9097 | 0.0033 |
| **Genotype:Sample Type** | 0.1964 | 0.0075 | 0.0864 | 0.0102 | 0.9172 | 0.0033 |
| **Sex:Genotype:Sample Type** | 0.1752 | 0.0078 | 0.1977 | 0.0079 | 0.9905 | 0.0019 |

**Supplemental Table 4. R2 and p values from mixed-effect PERMANOVAs for unweighted UniFrac distances**. P values < 0.05 shown in bold.

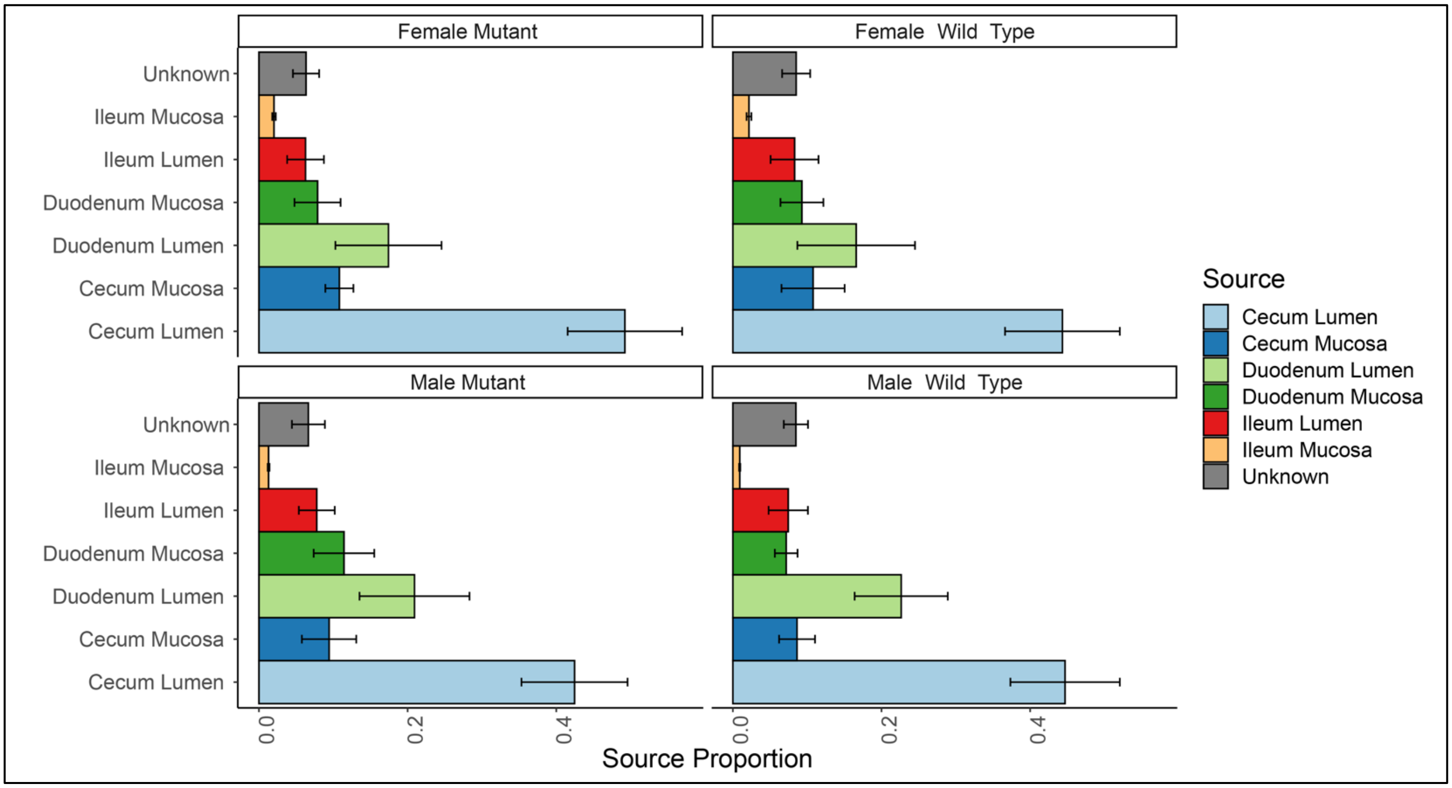

**Supplemental Figure 1 |Source contribution to feces by *hpg* genotype and sex**. 95% confidence intervals shown for SourceTracker source proportions of feces (sink) samples by sex and *hpg* genotype.

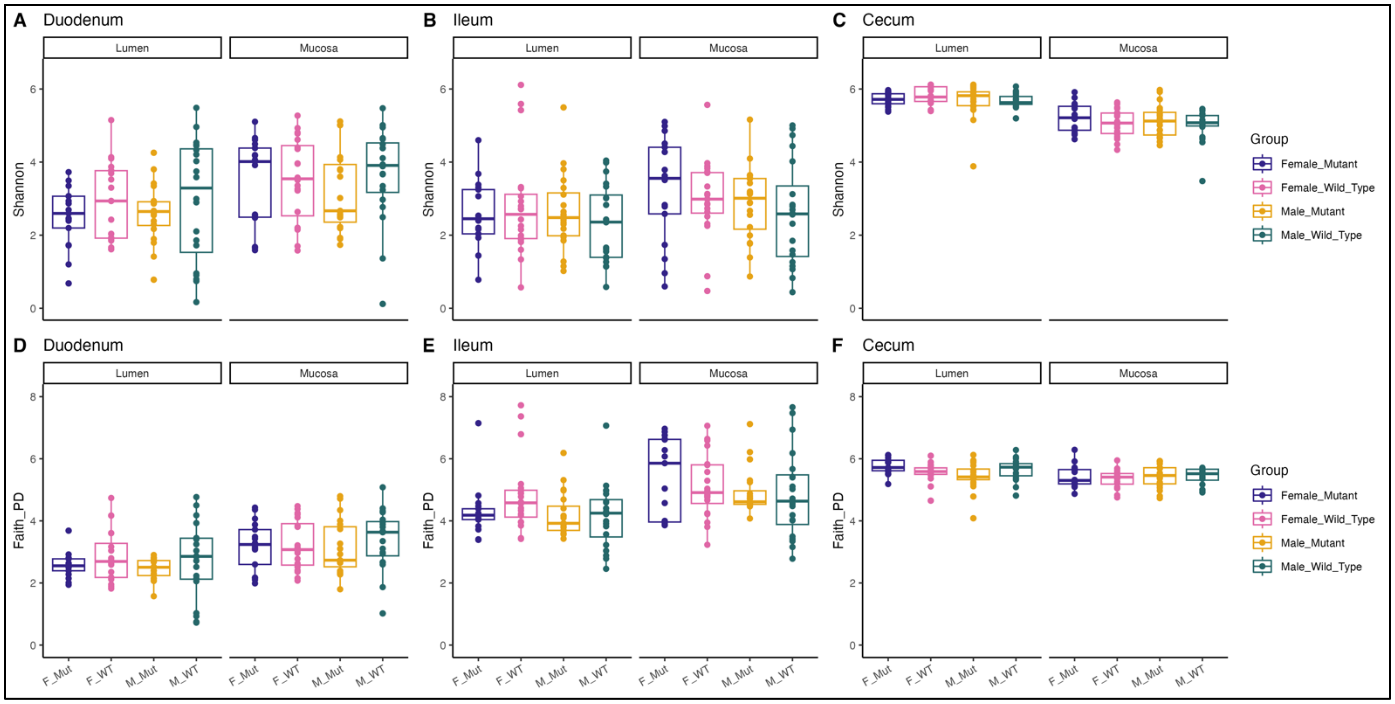

**Supplemental Figure 2 | Alpha diversity measures by sex and *hpg* genotype in each intestinal environment. A-C** Shannon index and **D-E** Faith's PD by sex and *hpg* genotype for the duodenum, ileum, and cecum.

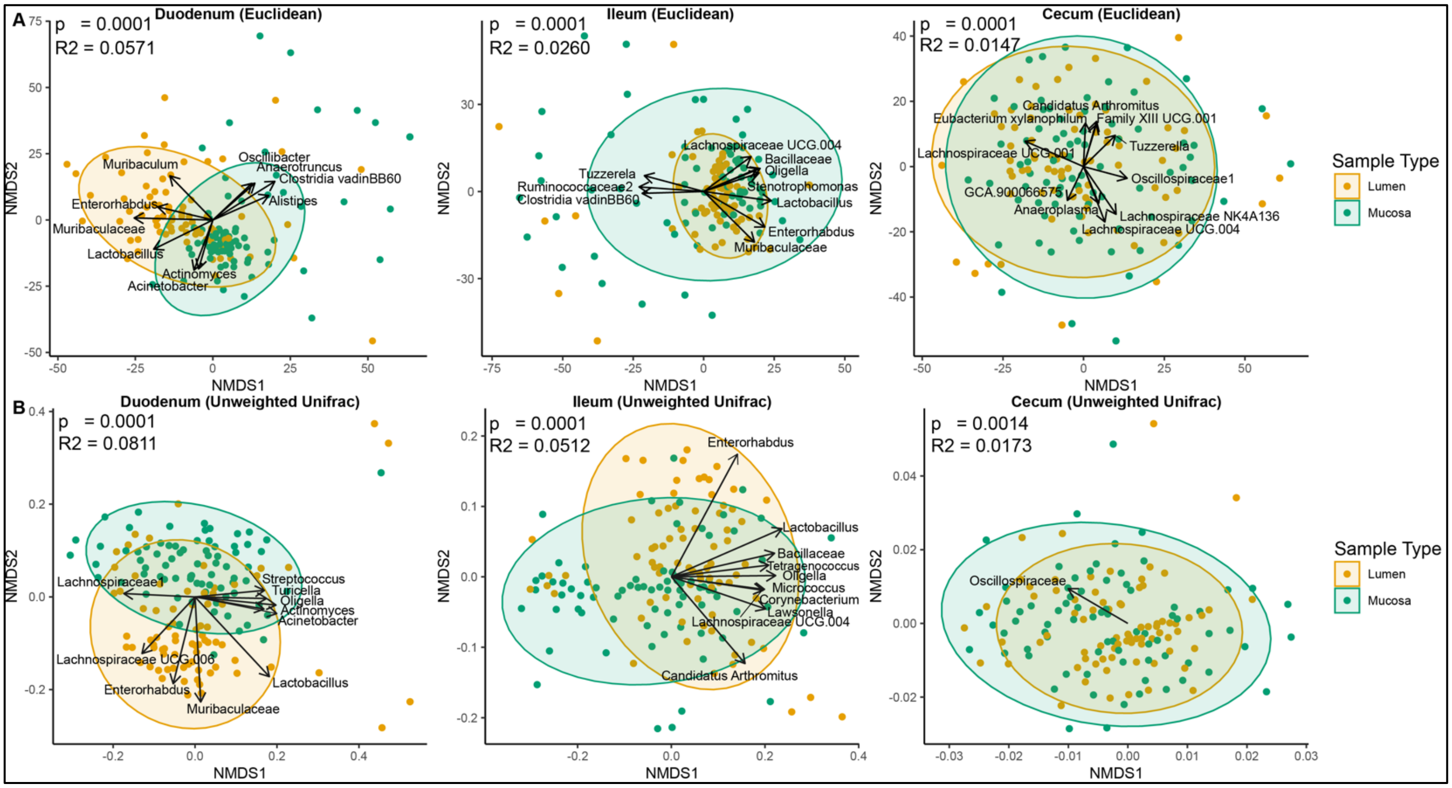

**Supplemental Figure 3. Beta diversity differences between lumen and mucosa**. **A** NMDS orientation plots of Euclidean distances comparing lumen and mucosa in the duodenum, ileum, and cecum. **B** NMDS plots of unweighted UniFrac distances for the duodenum, ileum, and cecum. The clr-transformed counts of genera were fit to each ordination and arrows are the vector average of the genus. Genera shown had the top 10 $R^2$ values of significant genera fit to the ordination (FDR-corrected p values < 0.05 determined by permutation test).

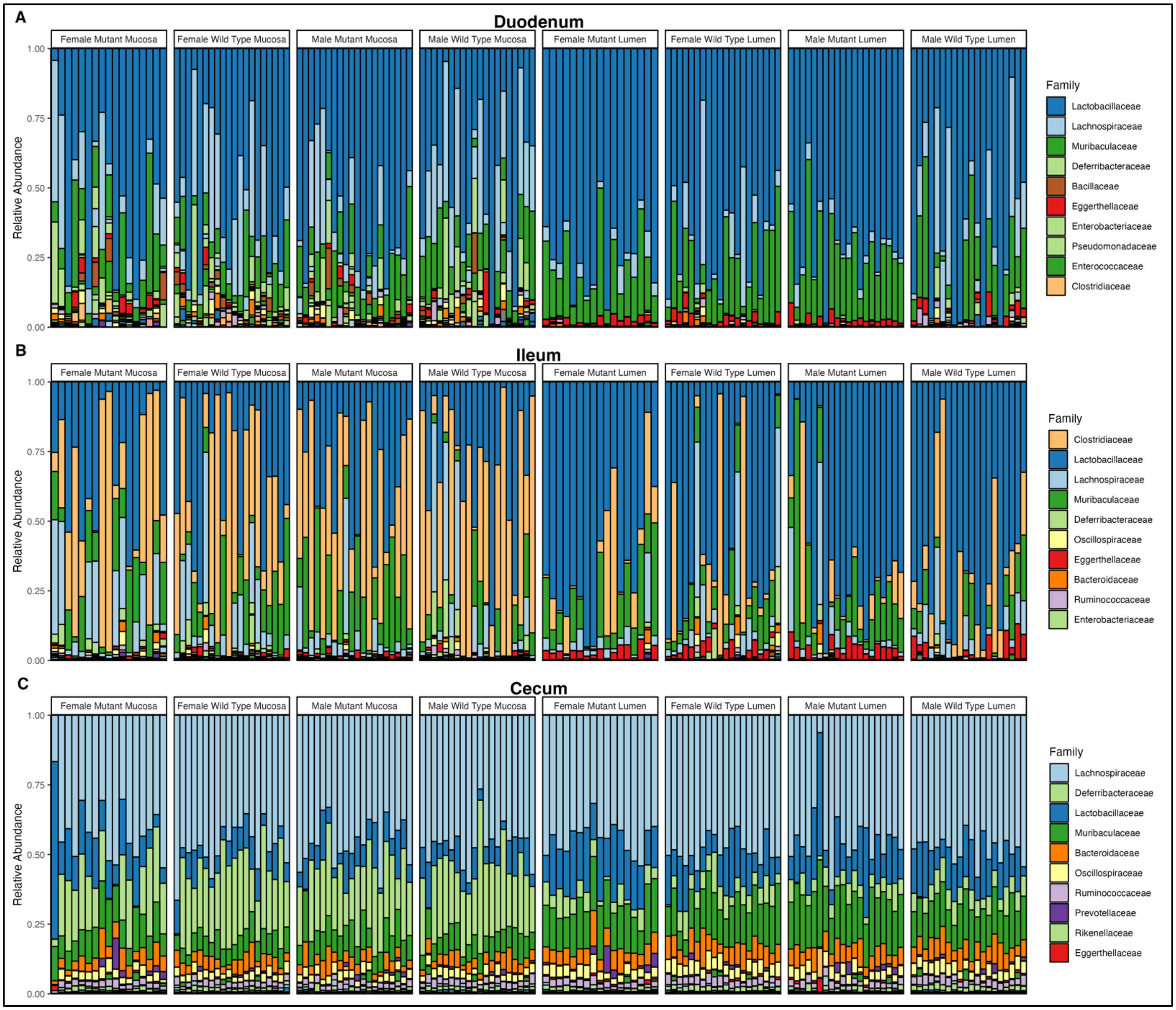

**Supplemental Figure 4 |Family relative abundance for each individual sample**. Relative abundance of each family for **A** duodenum, **B** ileum, and **C** cecum. Each bar represents a mouse intestinal sample.

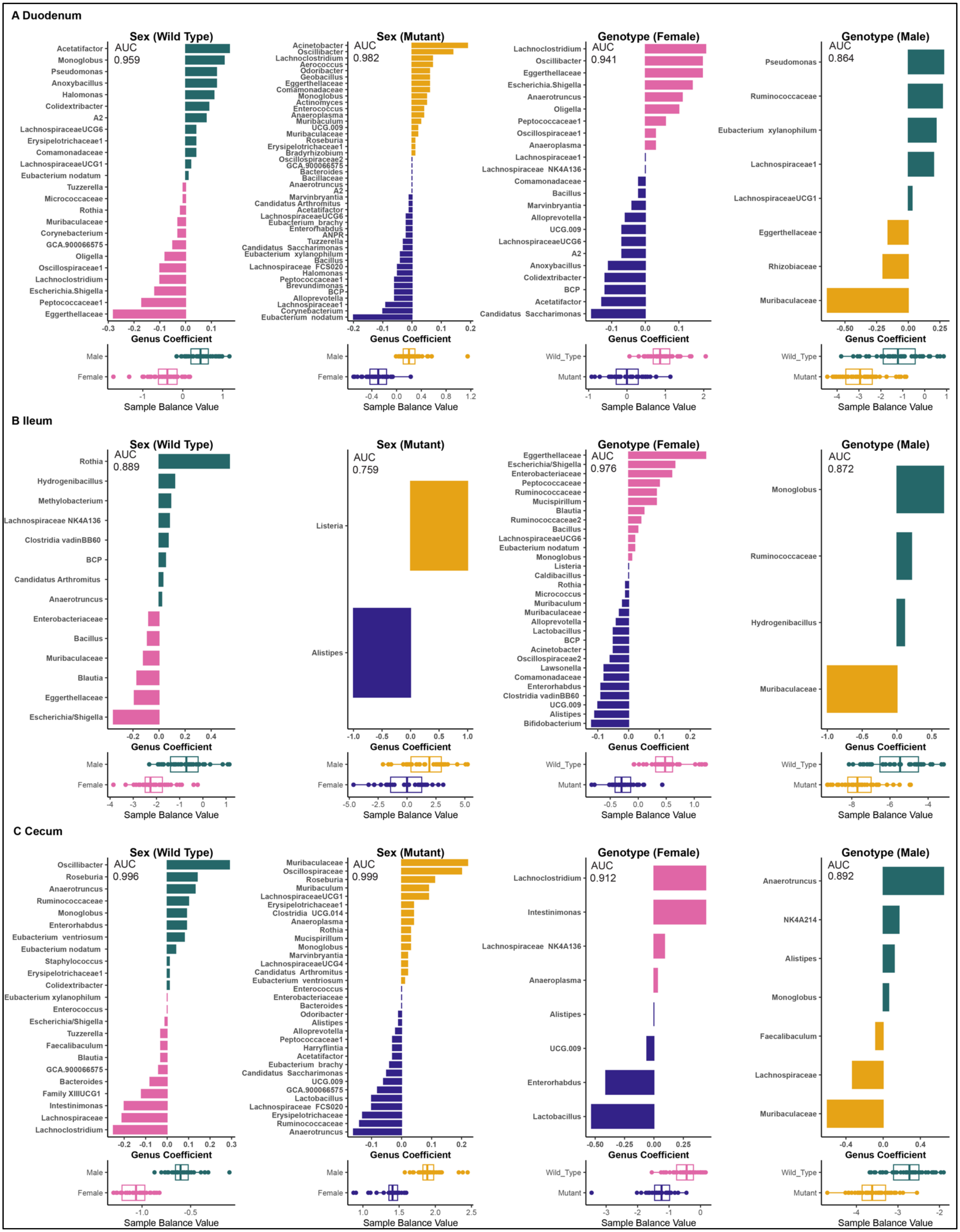
A Duodenum
Sex (Wild Type)
AUC
0.959
Acetatifactor
Monoglobus
Pseudomonas
Anoxybacillus
Halomonas
Colidextribacter
A2
LachnospiraceaeUCG6
Erysipelotrichaceae1
Comamonadaceae
LachnospiraceaeUCG1
Eubacterium nodatum
Tuzzerella
Micrococcaceae
Rothia
Muribaculaceae
Corynebacterium
GCA.900066575
Oligella
Oscillospiraceae1
Lachnoclostridium
Escherichia.Shigella
Peptococcaceae1
Eggerthellaceae
Genus Coefficient
Male
Female
Sample Balance Value
Sex (Mutant)
AUC
0.982
Acinetobacter
Oscillibacter
Lachnoclostridium
Aerococcus
Odoribacter
Geobacillus
Eggerthellaceae
Comamonadaceae
Monoglobus
Actinomyces
Enterococcus
Anaeroplasma
Muribaculum
UCG.009
Muribaculaceae
Roseburia
Erysipelotrichaceae1
Bradyrhizobium
Oscillospiraceae2
GCA.900066575
Bacteroides
Bacillaceae
Anaerotruncus
A2
Marvinbryantia
Candidatus Arthromitus
Acetatifactor
LachnospiraceaeUCG6
Eubacterium brachy
Enterorhabdus
ANPR
Tuzzerella
Candidatus Saccharimonas
Eubacterium xylanophilum
Bacillus
Lachnospiraceae FCS020
Halomonas
Peptococcaceae1
Brevundimonas
BCP
Alloprevotella
Lachnospiraceae1
Corynebacterium
Eubacterium nodatum
Genus Coefficient
Male
Female
Sample Balance Value
Genotype (Female)
AUC
0.941
Lachnoclostridium
Oscillibacter
Eggerthellaceae
Escherichia.Shigella
Anaerotruncus
Oligella
Peptococcaceae1
Oscillospiraceae1
Anaeroplasma
Lachnospiraceae1
Lachnospiraceae NK4A136
Comamonadaceae
Bacillus
Marvinbryantia
Alloprevotella
UCG.009
LachnospiraceaeUCG6
A2
Anoxybacillus
Colidextribacter
BCP
Acetatifactor
Candidatus Saccharimonas
Genus Coefficient
Wild_Type
Mutant
Sample Balance Value
Genotype (Male)
AUC
0.864
Pseudomonas
Ruminococcaceae
Eubacterium xylanophilum
Lachnospiraceae1
LachnospiraceaeUCG1
Eggerthellaceae
Rhizobiaceae
Muribaculaceae
Genus Coefficient
Wild_Type
Mutant
Sample Balance Value
B Ileum
Sex (Wild Type)
AUC
0.889
Rothia
Hydrogenibacillus
Methylobacterium
Lachnospiraceae NK4A136
Clostridia vadinBB60
BCP
Candidatus Arthromitus
Anaerotruncus
Enterobacteriaceae
Bacillus
Muribaculaceae
Blautia
Eggerthellaceae
Escherichia/Shigella
Genus Coefficient
Male
Female
Sample Balance Value
Sex (Mutant)
AUC
0.759
Listeria
Alistipes
Genus Coefficient
Male
Female
Sample Balance Value
Genotype (Female)
AUC
0.976
Eggerthellaceae
Escherichia/Shigella
Enterobacteriaceae
Peptococcaceae
Ruminococcaceae
Mucispirillum
Blautia
Ruminococcaceae2
Bacillus
LachnospiraceaeUCG6
Eubacterium nodatum
Monoglobus
Listeria
Caldibacillus
Rothia
Micrococcus
Muribaculum
Muribaculaceae
Alloprevotella
Lactobacillus
BCP
Acinetobacter
Oscillospiraceae2
Lawsonella
Comamonadaceae
Enterorhabdus
Clostridia vadinBB60
UCG.009
Alistipes
Bifidobacterium
Genus Coefficient
Wild_Type
Mutant
Sample Balance Value
Genotype (Male)
AUC
0.872
Monoglobus
Ruminococcaceae
Hydrogenibacillus
Muribaculaceae
Genus Coefficient
Wild_Type
Mutant
Sample Balance Value
C Cecum
Sex (Wild Type)
AUC
0.996
Oscillibacter
Roseburia
Anaerotruncus
Ruminococcaceae
Monoglobus
Enterorhabdus
Eubacterium ventriosum
Eubacterium nodatum
Staphylococcus
Erysipelotrichaceae1
Colidextribacter
Eubacterium xylanophilum
Enterococcus
Escherichia/Shigella
Tuzzerella
Faecalibaculum
Blautia
GCA.900066575
Bacteroides
Family XIIIUCG1
Intestinimonas
Lachnospiraceae
Lachnoclostridium
Genus Coefficient
Male
Female
Sample Balance Value
Sex (Mutant)
AUC
0.999
Muribaculaceae
Oscillospiraceae
Roseburia
Muribaculum
LachnospiraceaeUCG1
Erysipelotrichaceae1
Clostridia UCG.014
Anaeroplasma
Rothia
Mucispirillum
Monoglobus
Marvinbryantia
LachnospiraceaeUCG4
Candidatus Arthromitus
Eubacterium ventriosum
Enterococcus
Enterobacteriaceae
Bacteroides
Odoribacter
Alistipes
Alloprevotella
Peptococcaceae1
Harryflintia
Acetatifactor
Eubacterium brachy
Candidatus Saccharimonas
UCG.009
GCA.900066575
Lactobacillus
Lachnospiraceae FCS020
Erysipelotrichaceae
Ruminococcaceae
Anaerotruncus
Genus Coefficient
Male
Female
Sample Balance Value
Genotype (Female)
AUC
0.912
Lachnoclostridium
Intestinimonas
Lachnospiraceae NK4A136
Anaeroplasma
Alistipes
UCG.009
Enterorhabdus
Lactobacillus
Genus Coefficient
Wild_Type
Mutant
Sample Balance Value
Genotype (Male)
AUC
0.892
Anaerotruncus
NK4A214
Alistipes
Monoglobus
Faecalibaculum
Lachnospiraceae
Muribaculaceae
Genus Coefficient
Wild_Type
Mutant
Sample Balance Value

**Supplemental Figure 5 | Strength and complexity of genus-level balance models.** Sex differences in wild-type and mutant mice and *hpg* genotype differences in female and male mice in the **A** duodenum, **B** ileum, and **C** cecum. AUC (area under the curve) indicates the model strength (AUC of 1 indicates a model with perfect predictive value). Abbreviations: ANPR genus: *Allorhizobium /Neorhizobium/Pararhizobium/Rhizobium* genus group. BCP genus: *Burkholderia/Caballeronia/Paraburkholderi* genus group.

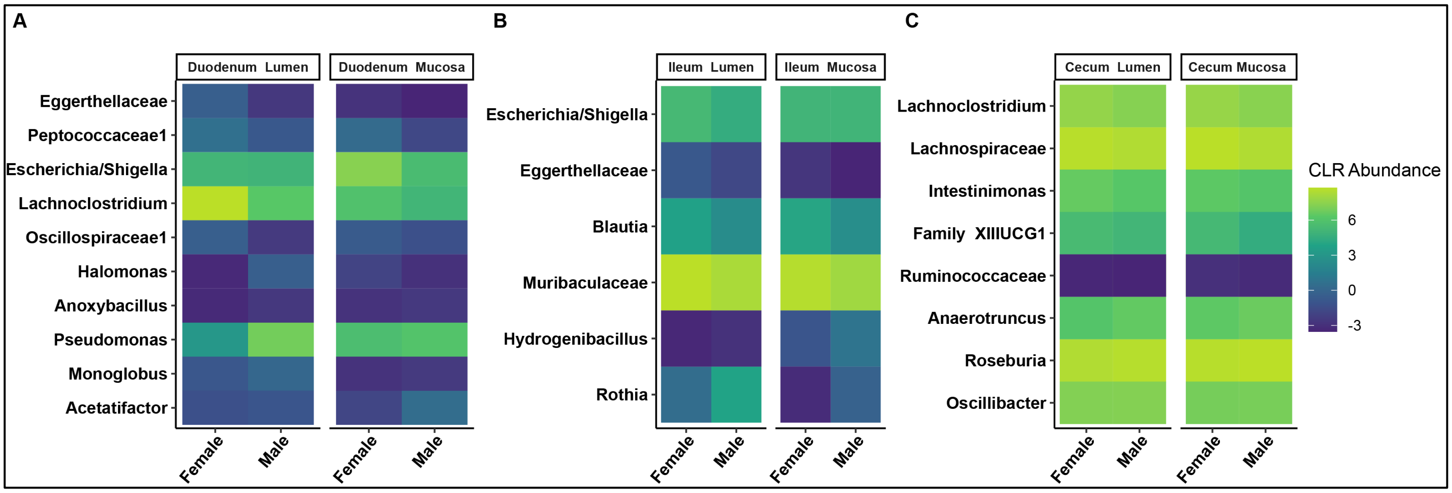

**Supplemental Figure 6 | Relative abundance of genera in *hpg* wild-type balance differs by sample type.** Heatmaps comparing genera abundances between wild-type sexes separating lumen and mucosa for the **A** duodenum, **B** ileum, and **C** cecum.
